# Local adaptation of *Legionella pneumophila* within a hospital hot water system increases tolerance to copper

**DOI:** 10.1101/2020.04.22.054569

**Authors:** Emilie Bédard, Hana Trigui, Jeffrey Liang, Margot Doberva, Kiran Paranjape, Cindy Lalancette, Sebastien P. Faucher, Michèle Prévost

## Abstract

In large-building water systems, *Legionella pneumophila* is exposed to common environmental stressors such as copper. The aim of this study was to evaluate the susceptibility to copper of *L. pneumophila* isolates recovered from various sites: two clinical and seven environmental from hot water systems biofilm & water, and from cooling tower water. After one-week acclimation in simulated drinking water, strains were exposed to various copper concentrations (0.8 to 5 mg/L) for over 672 hours. Complete loss of culturability was observed for three isolates, following copper exposure to 5 mg/L for 672h. Two ST1427-like isolates were highly sensitive to copper, while the other two, isolated from biofilm samples, were resistant. The expression of the copper resistance gene *copA* evaluated by RT-qPCR was significantly higher for the biofilm isolates. All four ST1427-like isolates were recovered from the same water system during an outbreak. Whole genome sequencing results confirmed that the four isolates are very close phylogenetically, differing by only 29 single nucleotide polymorphisms, suggesting *in situ* adaptation to microenvironmental conditions, possibly due to epigenetic regulation. These results indicate that the immediate environment within a building water distribution system influences the tolerance of *L. pneumophila* to copper. Increased contact of *L. pneumophila* biofilm strains with copper piping or copper alloys in the heat exchanger might lead to local adaptation. The phenotypic differences observed between water and biofilm isolates from the hot water system of a healthcare facility warrants further investigation to assess the relevance of evaluating disinfection performances based on water sampling alone.

**Importance:** *Legionella pneumophila* is a pathogen indigenous to natural and large building water systems in the bulk and the biofilm phases. The immediate environment within a system can impact the tolerance of *L. pneumophila* to environmental stressors, including copper. In healthcare facilities, copper levels in water can vary, depending on water quality, plumbing materials and age. This study evaluated the impact of the isolation site (water vs biofilm, hot water system vs cooling tower) within building water systems. Closely related strains isolated from a healthcare facility hot water system exhibited variable tolerance to copper stress shown by differential expression of *copA*, with biofilm isolates displaying highest expression and tolerance. Relying on the detection of *L. pneumophila* in water samples following exposure to environmental stressor such as copper may underestimate the prevalence of *L. pneumophila*, leading to inappropriate risk management strategies and increasing the risk of exposure for vulnerable patients.

## Introduction

*Legionella pneumophila* is an opportunistic pathogen that is the causative agent of Legionnaires’ disease and Pontiac fever (Fields et al., 2002). Between 2013 and 2014, 88% of hospitalizations and 100% of deaths resulting from drinking water-associated outbreaks in the United States were attributed to *Legionella* infections (Benedict et al., 2017). From 2000 to 2014, a 286% increase in legionellosis cases in the United States was reported, further highlighting the importance of this emerging pathogen (Garrison et al., 2016). Similarly Legionnaires’ disease cases have steadily increased in Europe between 2011 and 2016, with 81% of infections attributed to *L. pneumophila* serogroup 1 (European Centre for Disease Prevention and Control (ECDC), 2017). While the average mortality rate associated with legionellosis is estimated to be approximately 8% (Centers for Disease Control and Prevention (CDC), 2017; European Centre for Disease Prevention and Control (ECDC), 2017)), it can reach up to 25% in healthcare-associated outbreaks (Soda et al., 2017). The transmission of *L. pneumophila* to humans occurs through man-made water systems (Fields et al., 2002). Indeed, *Legionella* is known to proliferate in engineered water systems, such as cooling towers and large-building water distribution systems (Buse et al., 2012). In healthcare facilities, where the mortality rates are higher, hot water systems feeding taps and showers have a higher prevalence of *L. pneumophila* relative to other *Legionella* species (Bargellini et al., 2011; Boppe et al., 2016; Marchesi et al., 2011). Several factors inherent to large-building water systems are favorable to *L. pneumophila* persistence: lukewarm water temperature, presence of amoeba, use of disinfectants, stagnation, presence of biofilm, and plumbing materials (Buse et al., 2012; National Academies of Sciences, 2019).

The impact of plumbing material on microbial growth and development of biofilm is well-recognized (Proctor et al., 2016; Wang et al., 2014; Yu et al., 2010). Similar or increased incorporation and persistence of *Legionella* were measured by non-culture-based methods in copper grown biofilms compared to PVC, uPVC or PEX grown biofilm (Buse et al., 2014; Giao et al., 2015; Moritz et al., 2010). In contrast, a study reported a mean reduction of 2.5 log of *L. pneumophila* (measured by gene copies) in water within copper pipes compared to PEX pipes at temperatures ≤41°C, while total bacterial gene copies remained unchanged (Proctor et al., 2017). Because of its reported bactericidal effect, copper has long been used as a disinfecting material (Casey et al., 2010; Schmidt et al., 2012). Furthermore, copper-silver ionization treatment has been reported to reduce levels of *L. pneumophila* in hospital hot water distribution systems (Lin et al., 2011). Nevertheless, unreliable eradication of *L. pneumophila* following copper-silver ionization were linked to legionellosis cases (Bédard et al., 2016b; Blanc et al., 2005; Demirjian et al., 2015). Regulatory agencies recommend that copper levels in drinking water be maintained below 1.3 to 2 mg /L (Health Canada, 2014; United States Environmental Protection Agency (USEPA), 2009; World Health Organization (WHO), 2011). These guideline values were established for aesthetic reasons since reported health effects on healthy adults were only associated with levels exceeding 4 mg/L (Araya et al., 2003; Olivares et al., 2001). However, the state of California revised their Public Health Goal to 0.3 mg/L in 2008 (California Environmental Protection Agency et al., 2008) in response to reported adverse effects on infants and young children at levels below 1 mg/L (Stenhammar, 1999).

In healthcare facilities, copper levels in water carried within copper plumbing can vary between <10 μg/L and 800 μg/L, depending on water corrosiveness and pipe age (Bargellini et al., 2011; Bedard et al., 2015). These levels are even higher in new copper pipes, where dissolved copper can exceed 5 mg/L (Edwards et al., 2011). Reports of *L. pneumophila’s* ability to withstand copper stress are contradictory. *L. pneumophila* has been recovered from water within copper-pipe premise plumbing carrying up to 0.8 mg Cu/L (Bargellini et al., 2011; Boppe et al., 2016), and culturable *L. pneumophila* has been observed at levels up to 8 mg/L in a laboratory study (Jwanoswki et al., 2017). Copper has also been reported to inhibit the growth of *L. pneumophila* (Lin et al., 1996). These reported differences in copper effect on *L. pneumophila* could be due to the variable ability of each strain to tolerate copper, underlined by the genetic makeup of each strain or adaptive evolution of tolerance due to previous exposure. Indeed, *L. pneumophila* strains may behave differently depending on the environment from which they are isolated, and/or environmental conditions to which they have adapted. Bacteria can resist high copper concentration in water which can be mediated by the expression of specific genes. The only gene known to mediate copper resistance in *L. pneumophila* is the copper-translocating P_IB_-type ATPase *copA* (Kim et al., 2009; Trigui et al., 2013). *copA* is encoded on a 100 Kb mobile genetic element whose excision is regulated by the sRNA binding regulator Hfq (Trigui et al., 2013). Its episomal form is linked to higher expression of *copA* and, therefore, an increased tolerance to copper.

The main objectives of this study were to: 1) evaluate the impact of the isolation site (clinical *vs* environmental, water *vs* biofilm, hot water system *vs* cooling tower) of *L. pneumophila* strains on their tolerance to various levels of copper in drinking water over a one-month exposure time and 2) assess whether isolates recovered from different locations, but identified as closely related strains, exhibit variable tolerance to a copper stress through differential expression of *copA*.

## Results

The culturability of nine *L. pneumophila* strains (Table 1) was assessed over a 4-week (672 hours) exposure period to various copper concentration (0.8 to 5 mg/L). Strains were isolated from clinical or environmental samples. Environmental strains were recovered from water and biofilm of the hot water distribution systems in two healthcare facilities (HCF-A and HCF-B) and from water in two cooling towers (CT-C and CT-D). All strains isolated from the HCF-B hot water system (1427.B.F; 1427.W.HE; 1427.B.HE) and a patient (1427.C) were isolated following an outbreak and were found to be closely related by sequence-based typing (ST1427) and pulsed-field gel electrophoresis (Bédard et al., 2016b). No infections were associated with the environmental strains isolated from HCF-A and from CT-C. The environmental strain 62.W was isolated from a cooling tower (CT-D) following a large outbreak of *L. pneumophila* in Québec city (Levesque et al., 2014). Clinical strain 62.C was isolated from a patient during the outbreak associated to CT-D.

**Table 1:**
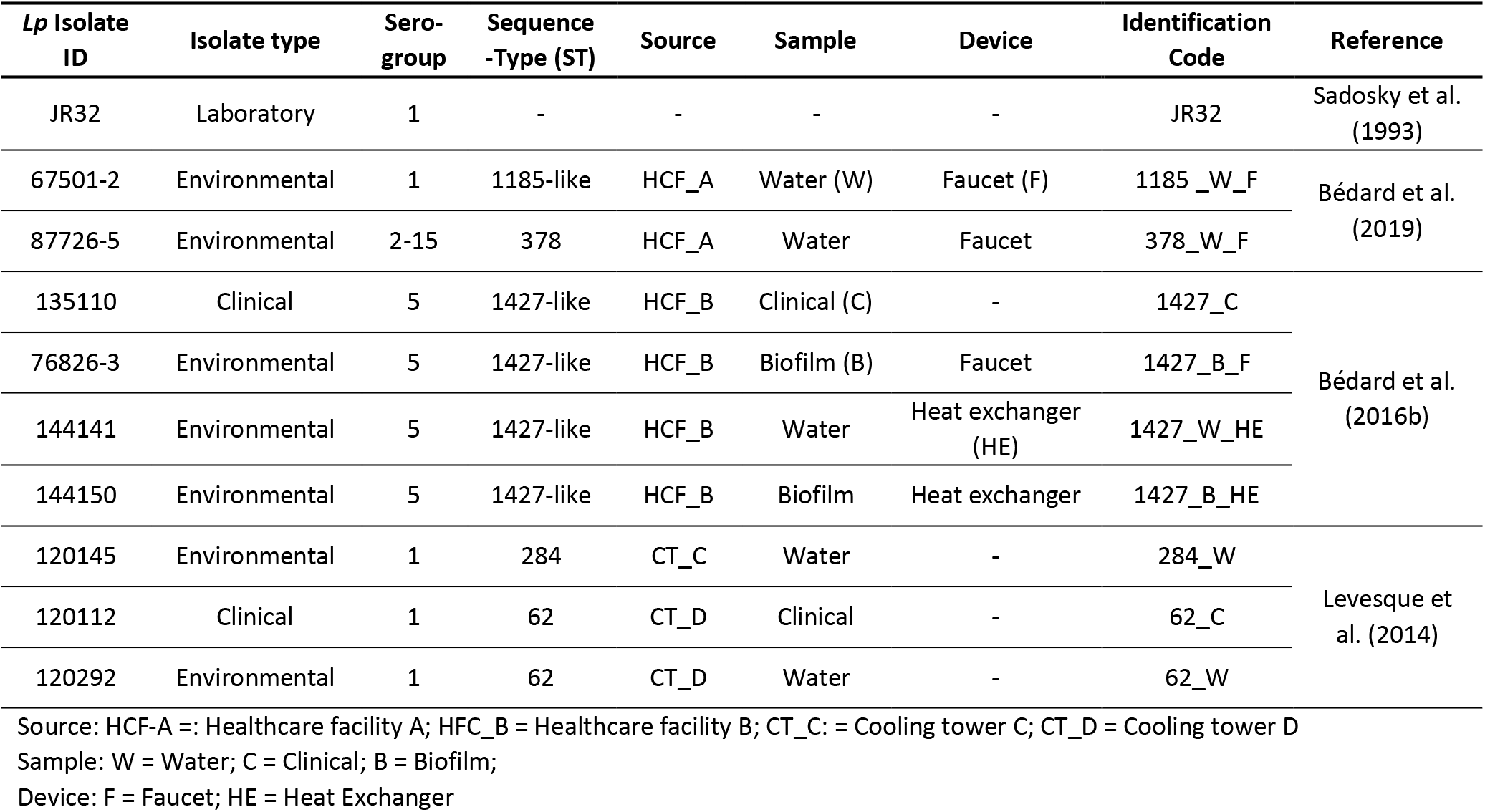
Characteristics of *L. pneumophila* isolates used in this study

Prior to copper exposure, strains were adapted to drinking water environment for one week, in simulated drinking water and at room temperature, to mimic conditions in premise plumbing. Following adaptation, the assay was conducted at 36°C, representative of water temperature in temperature-controlled faucets (electronic, foot operated or TMV) (Bédard et al., 2016d; Sydnor et al., 2012). In order to separate the decline of *L. pneumophila* culturability due to starvation and due to copper exposure, a control without copper and maintained at 36°C for the duration of the assay was monitored (Table S1). A general reduction in culturability was observed with increasing copper concentrations (Figure 1).

**Figure 1:**
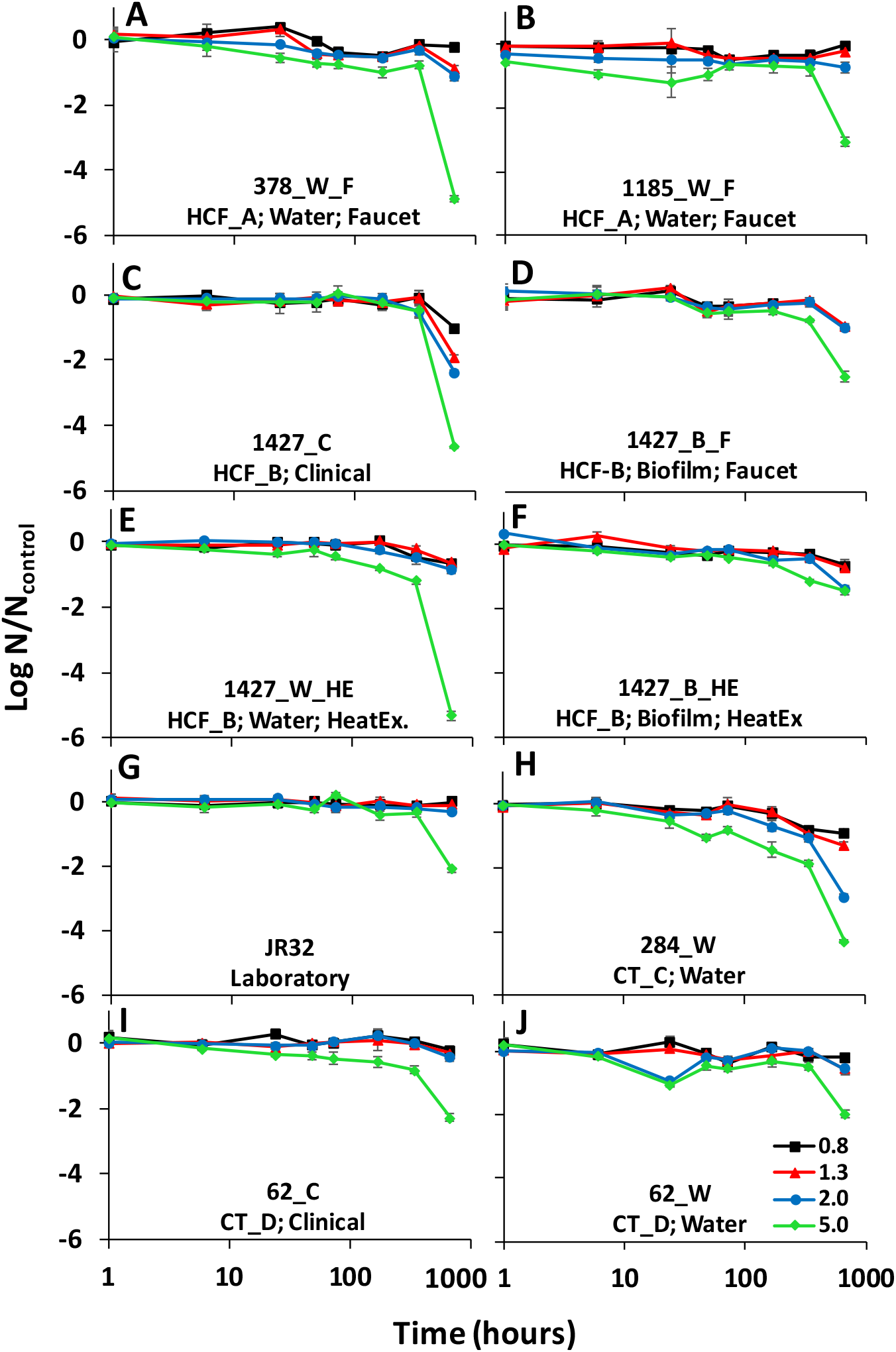
Survival of *L. pneumophila* isolates in copper-treated simulated drinking water relative to an untreated control in simulated drinking water. *L. pneumophila* strains were incubated with different copper concentrations (0.8; 1.3; 2; 5 mg /L) for a period of 1 to 672 hours. The log reduction in CFU counts was calculated relative to the untreated control for each strain. Results are grouped according to the origin of each isolate. (A-B) hot water system in healthcare facility A; (C-F) hot water system in healthcare facility B; G) laboratory strain JR32; H) cooling tower C; (I-J) cooling tower D. Error bars represent the standard deviation (n = 3).

There was no notable change at a concentration of 0.8 mg/L, with no more than 1 log reduction compared to the control after 672 hours. Similar results were obtained at a concentration of 1.3 mg/L, except for strain 1427.C, where 1.9 log reduction was observed after 672 hours. Partial loss of culturability was also observed after 672 hours at 2 mg/L copper concentrations for 1427.C and 284.W strains, with 2.4 and 2.9 log reduction respectively. The impact on culturability was clearly observed at 5 mg/L: most isolates showed a slow but steady decline over time, with a sharp drop in culturability between 336h and 672h (Figure 1). Copper tolerance varied greatly between strains after 672 hours of exposure (Figure 2). When comparing isolate pairs by a one-way ANOVA analysis, the log reduction was significantly different for 41 of the 45 pairs analyzed (Table S2). The four strains identified as belonging to ST1427 and isolated from the same healthcare facility clearly displayed different tolerance toward copper (Figure 2). Of note, 1427.B.HE was significantly more tolerant to copper then the other 1427-like strains, while 1427.B.F was more resistant then 1427.C and 1427.W.HE.

**Figure 2:**
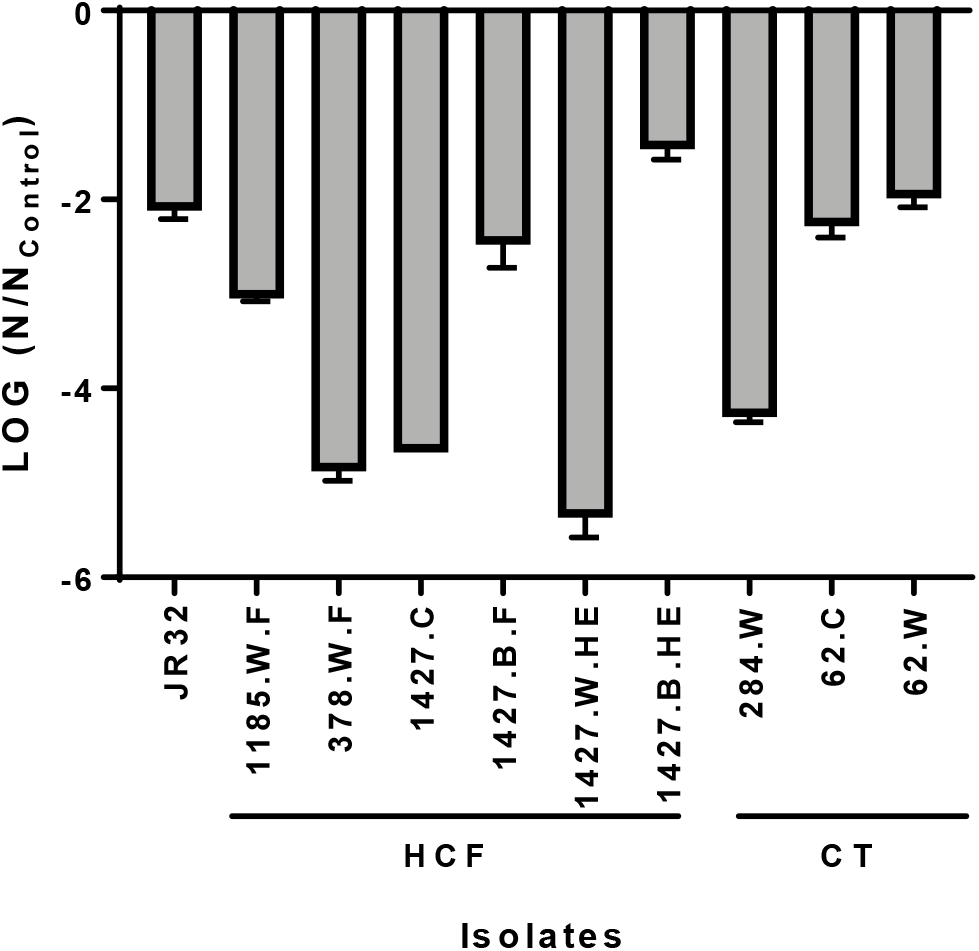
Log reduction of laboratory, environmental and clinical *L. pneumophila* isolates after exposure to 5 mg/L copper for 672 h in simulated drinking water. The log reduction in CFU counts is expressed as a ratio relative to the untreated control for each strain. Error bars represent the standard deviation (n = 3). CT : Cooling Tower and HCF : Health Care Facility.

Since the four strains belonging to the same ST displayed variable tolerance to copper, a more detailed investigation was conducted to test if the expression of *copA* gene was comparable between the isolates. RT-qPCR was used to test expression of *copA* after 30 minutes exposure to copper. Strains 1427.B.F and 1427.B.HE induced strong expression of *copA* after copper exposure, while 1427.HE and 1427.C did not (Figure 3). The expression of *copA* are in agreement with the resistance of the strains to copper exposure.

**Figure 3:**
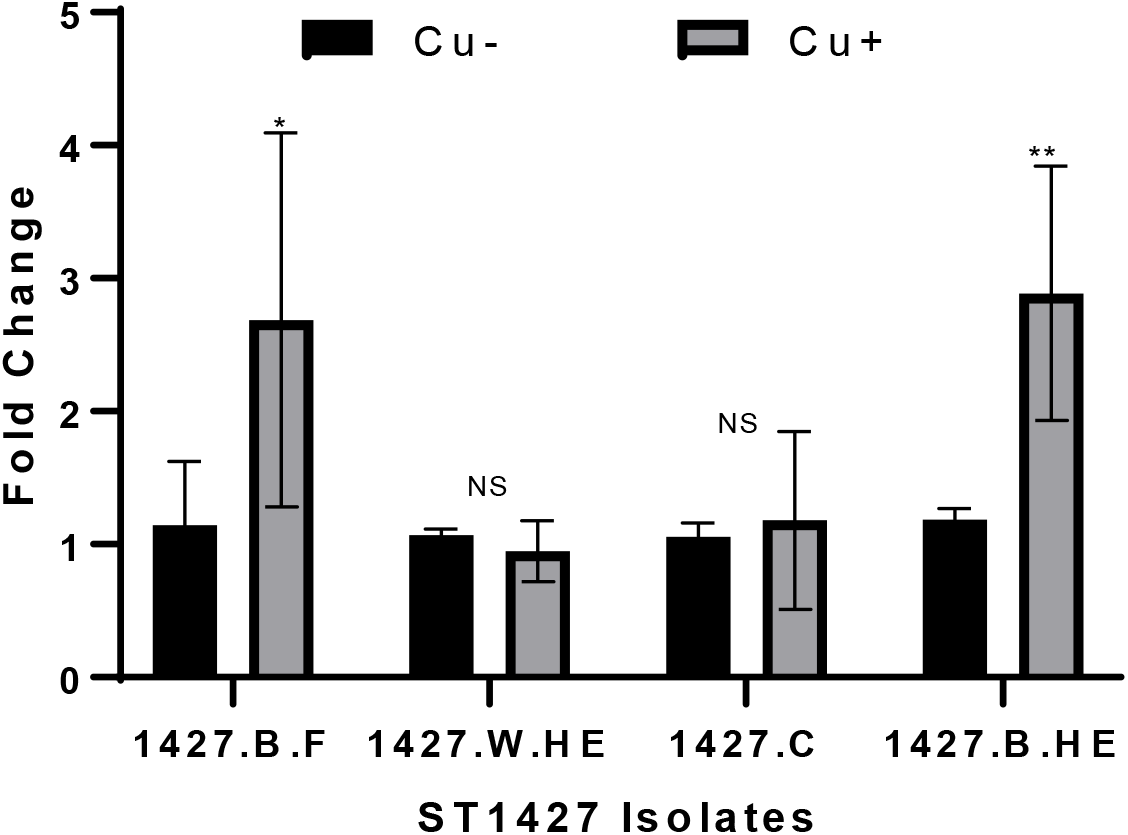
Expression of *copA* after 30-minute exposure to copper concentration of 5 mg/L in simulated drinking water at 36°C. RT-qPCR was used to test expression of *copA* in untreated (Cu −) and treated (Cu +) *L. pneumophila*. The fold change represents the change in expression after 30 minutes. Three replicates were used. We used an unpaired Student *t* test to assess statistical significance for each time point. *,*P* ≤ 0.05; **,*P* ≤ 0.001; (*versus* untreated).

Whole genome sequencing was conducted to investigate the phylogenetic relationship between the strains and identify genetic determinants of the phenotypic difference between the strains. The Microbial Genome Atlas was used to find the closest strains in the database (Rodriguez-R et al., 2018). Based on amino acid identity, 1427.B.F was more similar to *Lp* strain Thunder Bay (98.5% for 92.32% of protein shared) and to *Lp* strain Philadelphia 1 (98.4% for 94.11% proteins shared). This was supported by Mash (Ondov et al., 2016), with k-mers from contigs of the four isolates assembled by SPAdes (Bankevich et al., 2012) matching most closely to *Lp* Thunder Bay out of 96 full *Lp* genomes (Table S5).

A high-level phylogenetic tree was constructed using RAxML-ng (Kozlov et al. 2019) with SNPs situated outside of predicted recombinant regions called by Snippy and Gubbins (Seemann 2015, Croucher et al. 2015) to place the four isolates in a genomic context amongst 96 fully-completed *Lp* genomes (Figure S1). The four strains sequenced in this study formed a clade embedded in a cluster of Philadelphia-1 like strains. A higher-resolution tree was constructed using the same process to place the 1427-like isolates amongst closely-related strains (Figure 4), showing them to form a sister clade to a clade of strains rooted by *Lp* Thunder Bay (Khan 2013).

**Figure 4:**
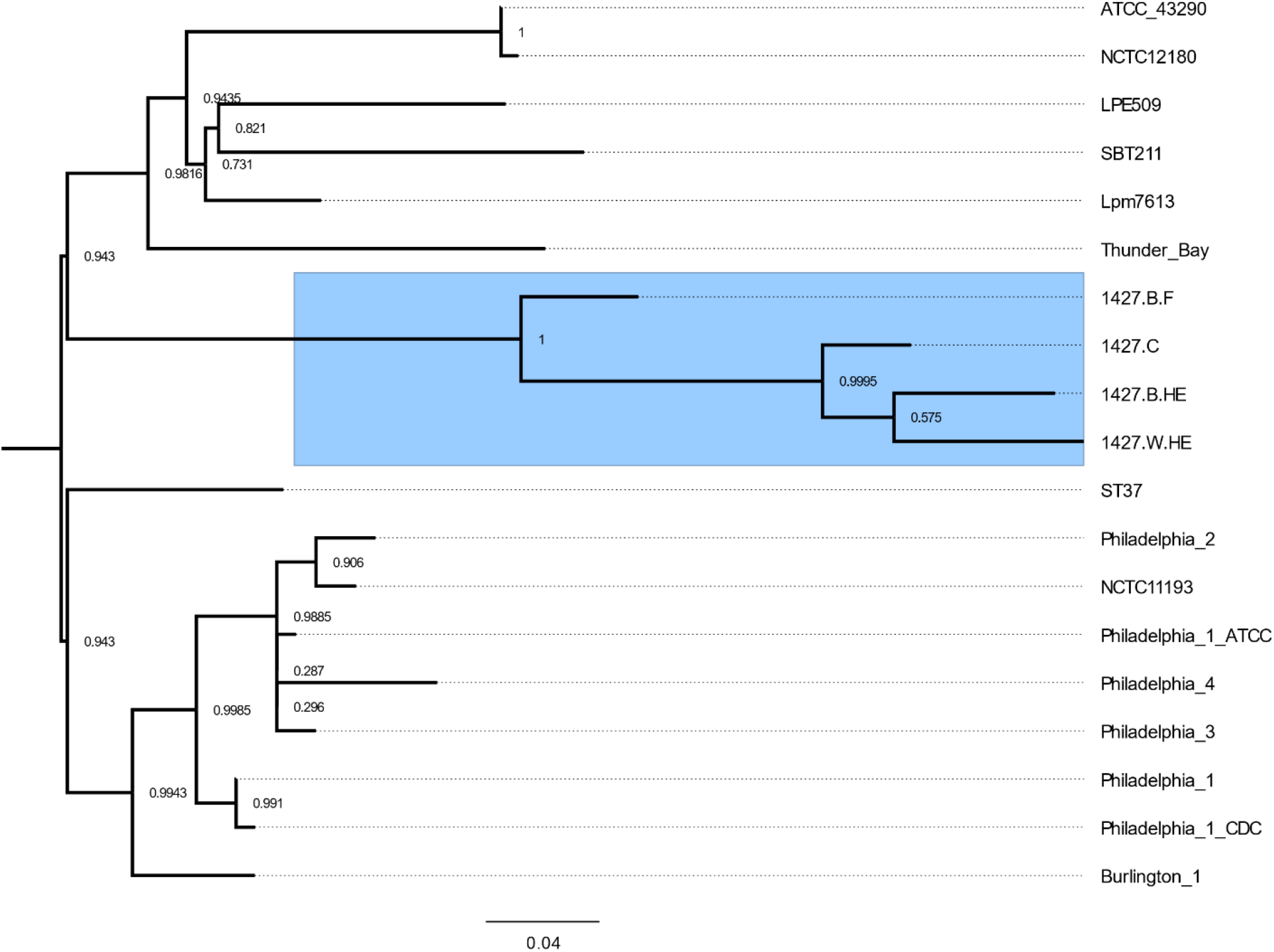
Whole genome-based phylogeny of the 1427-like isolates. 200 SNPs falling outside of recombinant regions were used to generate a phylogenetic tree of the 4 new isolates in the context of closely-related Philadelphia-like *Lp strains*. The four 1427-like isolates are highlighted in blue. Branch support was evaluated using 1000 bootstrap replicates in RAxML-ng transfer bootstrap expectation mode. Scale bar shows mean nucleotide substitutions per site.

The pangenome of the four 1427-like isolates was then analysed using ROARY (Page et al., 2015). Eight genes were identified as being present in only a subset of the isolates, but manual inspection showed these to be likely false negatives introduced during gene annotation. Only 29 SNPs were identified between the strains, confirming that the isolates are closely related (Table S3). Interestingly, the isolates from the heat exchanger (1427.B.HE, 1427.W.HE) were the most genetically similar to the clinical isolate (1427.C).

## Discussion

The tolerance against copper of nine *L. pneumophila* isolates recovered from environmental or clinical sources, in biofilm or water, belonging to various serogroup and sequence-types was evaluated in simulated drinking water. Experimental conditions were selected to best mimic premise plumbing environment. Firstly, cells were grown to the post-exponential phase, representative of the transmissive phase of *L. pneumophila*. In an engineered water system context, *L. pneumophila* would likely enter water after being expelled from a host cell, in the transmissive state. Post-exponential cells are also markedly more resistant to stress than exponential phase bacteria (Al-Bana et al., 2014; Byrne and Swanson, 1998; Hales and Shuman, 1999; Hammer et al., 2002). Secondly, cells were acclimatized in simulated drinking water at room temperature for 1 week prior to copper exposure. Starvation has been shown to increase the tolerance of *L. pneumophila* to numerous stresses (Bandyopadhyay et al., 2004; Byrne and Swanson, 1998; Chang et al., 2007; Li et al., 2014; Lynch et al., 2003). Lastly, following acclimation for 1 week in drinking water distribution system representative conditions, the tolerance to copper was tested in conditions simulating faucet temperatures, where a mix of cold and hot water is used and result in lukewarm temperature. For this reason, a temperature of 36°C was selected for the assay.

In this study, different *L. pneumophila* isolates displayed variable copper susceptibility over time when exposed to high concentrations of copper. The tolerance of all tested isolates to copper concentrations found in drinking water (≤ *2* mg/L) is not expected given the experimental conditions representative of drinking water plumbing. In contrast, distinct groups were observed at higher concentrations of 5 mg/L for prolonged contact time. The first group, including isolates from water and clinical sources (378.W.F, 1427.W.HE, 1427.C and 284.W), had lower tolerance to high copper doses and showed more than 4 log reduction after 672 hours of exposure. The second group included isolates from water, clinical and biofilm sources (JR32, 1185-W-F, 1427.B.F, 62.C and 62.W) and displayed 2 to 3 log reduction after 672 hours at 5 mg/L. Finally, the isolate 1427.B.HE was considered resistant to copper stress, as it showed less than 2 log reduction in culturability after 672 hours. No trend was observed suggesting that copper susceptibility is linked to sequence-type, although the number of strains from each ST was not sufficient to make any conclusion. While two isolates (patient-derived 62.C and cooling water 62.W) presented similar copper tolerance profiles at all tested concentrations (Figure 2), the four ST1427 strains had different tolerance profiles. In both cases, isolates were recovered during the investigation of *L. pneumophila* outbreaks (Bédard et al., 2016b; Levesque et al., 2016). When comparing a strain isolated from cooling tower water unrelated to an outbreak (284.W, ST284), the outbreak isolate displayed more resistance toward copper (Figure 2). As previously tested, the outbreak-associated strain (62.W) was significantly more infectious toward human macrophages, whereas the infectivity toward *Acanthamoeba castellanii* was comparable (Levesque et al., 2014).

The nature of the isolation site (environmental *vs* clinical, hot water system *vs* cooling towers, biofilm *vs* water) is also an important factor to consider when evaluating the copper resistance of environmental strains. The composition of drinking water and plumbing material have been shown to impact downstream microbial communities and the integration of *L. pneumophila* into biofilm (Lu et al., 2014). In this study, four isolates were recovered during an *L. pneumophila* outbreak investigation in a healthcare facility (HCF-B, Bédard et al. (2016b)). The source of the outbreak was identified as the hot water system, from which isolates 1427.B.F, 1427.W.HE and 1427.B.HE were recovered. These environmental isolates belong to the same sequence type (ST1427) (Bédard et al. (2016b)). In this study, whole genome sequencing analysis was performed on these four strains which confirm that they are closely related. Despite genomic similarities, we report that tolerance to copper stress was highly variable between the four isolates (Figure 2).

Microevolution within the population of *L. pneumophila* within hospitals water distribution systems was previously studied by whole genome sequencing (David et al., 2017). The strains isolated from a single hospital generally clustered according to the location. For examples, isolates from the same wards clustered together. However, all isolates from one hospital were related and most likely originated from a single ancestor. Therefore, it seems that water systems are seeded by one or a few strains that colonize the system and evolve within it, generating some limited diversity over time (David et al., 2017). Our results are in agreement with this hypothesis. Presumably, the water system was colonized initially by one 1427-like strain, which evolved within it over time, generating isolates with different phenotype. The number of SNPs identified between these strains are in the range of SNPs found between isolates from the same water systems (Bartley et al., 2016; David et al., 2017). Evolution of 1427-like population was driven by the persistence of the most adapted variants in the different microenvironment present in the water distribution systems, such as the tap or heat exchanger. Isolates 1427.B.HE and 1427.B.F displayed response to the presence of copper by significantly inducing higher expression of *copA* (Figure 3). Those two strains were recovered from environmental biofilm growing on metallic surfaces: stainless steel heat-exchanger plates containing chromium and nickel, and internal faucet surface. The ability of 1427.B.HE and 1427.B.F to maintain culturability in the presence of copper could be linked to their ability to form biofilm and their previous exposure to heavy metals from the substrata. Indeed, biofilm-derived cells have an increased resistance to heavy metals compared to suspended cells (Harrison et al., 2007). In the present case, the exposure of *L. pneumophila* to chrome and nickel during stagnation within the heat exchanger may have triggered integration into the biofilm, thereby activating metal resistance mechanisms, including those against copper (Baker-Austin et al., 2006). For example, the *L. pneumophila* gold-induced genes (GIG) operon is induced following exposure to copper, but the functional importance of this locus for copper resistance is unknown (Jwanoswki et al., 2017). The reported increased tolerance of drinking water bacteria vs. raw water bacteria to copper, lead and zinc further highlights the critical role of the environment in acquiring resistance (Calomiris et al., 1984).

The differences in copper resistance observed between the strains tested in this study (Figure 1) may be caused by differential *copA* expression as shown in Figure 2. However, resistance could also be associated to the presence of other, yet unidentified copper resistance genes. Unfortunately, no SNPs were found exclusively in the two most resistant strains. In addition, no non-synonymous mutation was found in genes associated with copper resistance. Therefore, SNPs or gene content cannot explain the difference in copper resistance and the difference in the expression of *copA* seen between the isolates, suggesting that these phenotypic differences are mediated by an epigenetic mechanism. Epigenetic regulation in bacteria involves DNA methylation, which affects the binding of transcriptional regulator to promoters (Sánchez-Romero and Casadesús, 2020). Epigenetic regulation can change gene expression without the need for DNA mutation. This process is mediated by DNA methyltransferase such as Dam (Sánchez-Romero and Casadesús, 2020). The role of DNA-methylation in gene regulation in bacteria is still poorly understood and has not been described yet in *L. pneumophila*. Nevertheless, evidence of Dam-mediated methylation of *Lp* genomic DNA was reported in 1996 (Lema and Brown, 1996). To our knowledge, this is the only study of DNA-methylation in *Lp*. In addition, DNA-methyltransferase are encoded in *Lp* genomes. *copA* is transcribed together with the upstream gene (lpg1034) from a transcription start site located at position −97 (Sahr et al., 2012). However, the promoter region does not contain GATC Dam methylation motifs, suggesting that a transcriptional regulator of *copA* might be under epigenetic regulation. Nevertheless, epigenetic regulation of *copA* is consistent with the transcriptional analysis showing that the resistant strains are responsive to the presence of copper, but the susceptible strains are not. Further study of epigenetic regulation in *Lp* will be required to investigate this possibility, which is outside the scope of this report.

These results suggest that the immediate environment can have a significant impact on the copper tolerance of closely related isolates. Our results suggest that broad susceptibility range in *L. pneumophila* isolates can be expected for other disinfectants. This, in turn, is critical information to consider when establishing disinfection strategies within a large building and selecting a sampling strategy for the monitoring of disinfection performance for *L. pneumophila* control. The selection of sampling sites within a given building’s water system and, sampling of water *vs*. biofilm may affect the evaluation of prevalence, and therefore, impact the subsequent risk assessment. The higher resistance of the biofilm-derived strains relative to suspended cells supports the overwhelming role of biofilm in promoting metal resistance. Importantly, an increase in resistance to metals is known to simultaneously lead to increased antibiotic resistance (Baker-Austin et al., 2006; Harrison et al., 2007). The difference observed between water and biofilm isolates in a healthcare facility hot water system questions the ability to evaluate disinfection performance based on water sampling alone and *L. pneumophila* isolated therein. An underestimation of *L. pneumophila* prevalence, combined with the exposure of vulnerable patients may lead to inappropriate risk management strategies.

## Material and methods

### Bacterial strains and culture conditions

Experiments were performed using nine *L. pneumophila* strains isolated from clinical or environmental samples, and *L. pneumophila* JR32, a Philadelphia-1 laboratory-adapted strain resistant to streptomycin (Sadosky et al., 1993). A summary of isolates used in this study and their characteristics is presented in Table 1. The clinical strains (1427.C & 62.C) and environmental strains (1427.W.HE, 1427.B.HE, 284.W & 62.W) were provided by the Laboratoire de Santé Publique du Québec. The average copper concentration in water from HCF-A and HCF-B were 0.57 mg/L and 0.40 mg/L respectively (Bédard et al., 2016a; Bedard et al., 2015). Isolates 1185-W-F, 1427.B.F, 378.W.F, 1427.W.HE, 1427.C and 1427.B.HE were typed by sequence-based typing, according to the European Working Group for *Legionella* Infections (EWGLI) protocol (Gaia et al., 2005; Ratzow et al., 2007). Sanger sequencing results were analyzed and a sequence type (ST) was assigned according to the EWGLI database.

Strains stored in 60% glycerol at −80°C were grown on Buffered Charcoal Yeast Extract (BCYE, Oxoid) agar for 3 days at 36°C. Resulting colonies were inoculated overnight at 36°C into sterile yeast extract broth (pH 6.88) with the growth supplement SR0110 (Oxoid). Cells were harvested by centrifugation (3000 g for 30 min), washed twice with sterile synthetic water and suspended in the same medium to an initial concentration of 10^8^ cells/mL. Prior to copper exposure, cells were starved in sterile synthetic water for 1 week at room temperature (22°C), to simulate environmental conditions in drinking water municipal distribution system and building main distribution pipes.

### Synthetic water preparation

Synthetic water was prepared by adding salts (final salt concentrations found in Table S4) to 0.22 μm filtered and autoclaved Milli-Q water. The water was buffered with 3-(N-Morpholino) propane sulfonic acid (MOPS, Sigma-Aldrich) to a final concentration of 1mM. The salts and their respective concentrations were selected to mimic Montreal tap water as per the Annual Drinking Water Quality Report (Ville de Montréal, 2015). The synthetic water was adjusted to an alkalinity of 60 mg CaCO_3_/L and a pH of 7.8. Individual stock salt solutions were prepared in sterile ultra-pure water. The MOPS buffer was selected due to its target pH range, low complexation rate with metal ions and since it does not interfere with cell growth (Ferreira et al., 2015). Final pH was adjusted to 7.8 using 0.1M NaOH.

### Experimental conditions

Each strain was starved in synthetic water for 1 week and subsequently, diluted in 20 mL of sterile synthetic water (final concentration of approximately 10^8^ cells/mL) with the following copper concentrations: 0, 0.8, 1.3, 2 and 5 mg/L. Copper chloride was selected as the copper source since it has the highest dissolved-to-total copper ratio in water (71 – 98% for copper chloride, compared to 51 – 67% for copper sulfate and 63% for copper nitrate). A high dissolved-to-total ion ratio was used to represent the behaviour of copper in drinking water, which is found mostly in the dissolved form. As an example in HCF-A, the mean dissolved-to-total ratio was 98% (n=75, 87 to 100%). Exposure to copper was carried out at 36°C to simulate temperature at the tap when hot and cold water are mixed. The survival of *L. pneumophila* isolates exposed to copper was monitored over time (1h, 6h, 24h, 48h, 72h, 168h, 336h and 672h) by culture. Water without a bacterial inoculum was used as a negative control. For each condition and time point, cell culturability was evaluated in triplicates on BCYE agar.

### copA expression

Analysis of *copA* gene expression was performed by reverse transcription-quantitative PCR (RT-qPCR). The isolates 1427.C, 1427.B.F, 1427.W.HE and 1427.B.HE were starved in synthetic drinking water (Table S4). Cells were collected prior and after exposure to 5 mg/L of copper for 30 min. RNA was extracted with TRIzol reagent according to the manufacturer’s protocol (Invitrogen). One microgram of RNA was then converted to cDNA by using random nonamers and ProtoScript II Reverse Transcriptase (New England Biolabs) according to the manufacturer’s instructions. For each sample, a negative control without reverse transcriptase was used. qPCRs were then performed with 1 μl of cDNA using Universal SYBR green Supermix (BIO-RAD). Primer sets used for real-time PCR analysis are qcopA forward (CAATACCCTGGTGGTCGATAAAAC) and qcopA reverse (TGCCGCAGCTAATGCTAAAG) (Trigui et al., 2013). A relative quantification strategy was used to perform analysis of the qPCR data. Transcript levels were normalized to 16S rRNA in each sample, using primers 16S_QF and 16S_QR (Trigui et al., 2015), and changes in the gene expression data from prior copper exposure to after exposure were calculated as fold change (Livak and Schmittgen, 2001). Statistical analysis between the strains exposed to and the wild-type control was performed using an unpaired one-tailed Student’s t-test.

### Whole Genome Sequencing of *Legionella pneumophila* strains

DNA libraries for sequencing on the MiSeq platform were made using the Nextera XT DNA library prep kit (Illumina). Briefly, the Wizard genomic DNA purification kit (Promega) was used to extract and purify the DNA from four *Legionella pneumophila* isolates, 1427_B_F, 1427_W_HE, 1427_C, and 1427_B_HE (Table 1). DNA quality and concentration were evaluated using a 0.8% agarose gel and the Quant-iT PicoGreen dsDNA assay kit (Thermofisher). Subsequent to the quality check, the genomic samples were prepared following Nextera XT manufacturer’s instructions. To ensure proper fragmentation of the genomic DNA, the DNA samples were evaluated on an Agilent Technology 2100 Bioanalyzer (Agilent). Once the DNA passed this quality control measure, the DNA samples were diluted to 2nM and pooled together to create the DNA library for sequencing. The library was frozen until further use (no more than 3 days). On the day of the sequencing run, 0.2 N NaOH was used to denature the DNA library. This denatured library was diluted to 12 pM loading concentration with HT1 buffer (Illumina). PhiX control (20 pM) was spiked at 1% into the library. The library was sequenced on an Illumina MiSeq platform using a V3 MiSeq Reagent kit (600 cycles). The raw reads of this sequencing project have been uploaded to the NCBI SRA database under the bioproject number PRJNA610262.

### WGS Read Analysis and Phylogeny Construction

Reads were pre-processed for quality control and adapter trimming using fastp 0.20.0 (Chen et al., 2018). SPAdes 3.13.1 (Bankevich et al., 2012) was used to assemble each of the four isolates into draft collections of contigs. These were compared using Mash 2.2 (Ondov et al., 2016) to all (96, as of January 22, 2020, Table S5) complete *L. pneumophila* genomes available through RefSeq. The four isolates were shown to form a clade which was most like strain Thunder Bay.

Analysis was conducted using Thunder Bay (NC_021350) as a reference genome for mapping and annotation. The snippy 4.4.5 (Seemann, 2014) pipeline (Garrison and Marth, 2012; Li, 2013) was used to call variant sites in the four isolates. Gubbins 2.4.1 (Croucher et al., 2014) was used to filter out probable recombinant sites from the SNP alignments. Phylogenies were constructed from the remaining SNP sites using RaxML-NG 0.9.0 (Kozlov et al., 2019) with model GTR+Gamma. To increase resolution, the same method was used to produce a phylogenetic tree using a subset of closely-related genomes (Table S5). Branch support was estimated using 1000 bootstrap replicates in transfer bootstrap expectation mode. Finally, Prokka 1.14.3 (Seemann, 2014) was used to annotate the genomes of the four isolates, and pan-genome analysis was conducted using Roary 3.12.0 (Page et al., 2015).

## Acknowledgements

This study was supported by the partners of the NSERC Industrial Chair on Drinking Water. The authors would like to thank Chair staff especially Jacinthe Mailly and Mélanie Rivard, Marie-Ève Benoit and Wendy Andriantsarafara. This work was supported by a NSERC Discovery Grant (RGPIN/04499-2018) to SPF. Jeffrey Liang is the recipient of an Alexander Graham Bell Canada Graduate Scolarships – Master’s.

